# Reproducible and accessible analysis of transposon insertion data at scale

**DOI:** 10.1101/2020.05.19.105429

**Authors:** Delphine Larivière, Laura Wickham, Kenneth C. Keiler, Anton Nekrutenko

## Abstract

Significant progress has been made in advancing and standardizing tools for human genomic and biomedical research, yet the field of next generation sequencing (NGS) analysis for microorganisms (including multiple pathogens) remains fragmented, lacks accessible and reusable tools, is hindered by local computational resource limitations, and does not offer widely accepted standards. One of such “problem areas” is the analysis of Transposon Insertion Sequencing (TIS) data. TIS allows perturbing the entire genome of a microorganism by introducing random insertions of transposon-derived constructs. The impact of the insertions on the survival and growth provides precise information about genes affecting specific phenotypic characteristics. A wide array of tools has been developed to analyze TIS data and among the variety of options available, it is often difficult to identify which one can provide a reliable and reproducible analysis. Here we sought to understand the challenges and propose reliable practices for the analysis of TIS experiments. Using data from two recent TIS studies we have developed a series of workflows that include multiple tools for data de-multiplexing, promoter sequence identification, transposon flank alignment, and read count repartition across the genome. Particular attention was paid to quality control procedures such as the determination of the optimal tool parameters for the analysis and removal of contamination. Our work provides an assessment of the currently available tools for TIS data analysis and offers ready to use workflows that can be invoked by anyone in the world using our public Galaxy platform (https://usegalaxy.org). To lower the entry barriers we have also developed interactive tutorials explaining details of TIS data analysis procedures at https://bit.ly/gxy-tis.

**Importance:** A wide array of tools has been developed to analyze TIS data and among the variety of options available, it is often difficult to identify which one can provide a reliable and reproducible analysis. Here we sought to understand the challenges and propose reliable practices for the analysis of TIS experiments. Using data from two recent TIS studies we have developed a series of workflows that include multiple tools for data de-multiplexing, promoter sequence identification, transposon flank alignment, and read count repartition across the genome. Particular attention was paid to quality control procedures such as the determination of the optimal tool parameters for the analysis and removal of contamination. Our work democratizes the TIS data analysis by providing open workflows supported by public computational infrastructure.

## Introduction

Transposition insertion sequencing (TIS) is based on random integration of transposons throughout a genome. These insertion knock out or alter expression of genes and functional elements. A TIS library—a population of bacterial cells carrying transposon insertions—is divided into aliquotes subjected to different experimental conditions. Upon completion of an experiment sites flanking transposon insertions are amplified and amplification products are subjected to high throughput sequencing (HTS). Mapping of resulting sequencing reads against the host genome reveals locations of insertions. Regions containing insertion can tolerate disruptions and thus are *non-essential*, while location void of insertions are likely under purifying selection and are *essential*. This approach led to successful genome-wide identification of essential and non-essential genes in a number of species^1^. A more comprehensive application of TIS involves insertion of transposon constructs carrying regulatory elements such as promoters^2^. In addition to binary readout (essential/non-essential) this approach yields information about effects of up- and down-regulation of specific genes.

Randomly pooled transposons libraries are commonly created with Mariner or Tn5 transposons. Mariner transposons target `TA` dinucleotides. They have wide species specificity and are stable^3^. The methods based on Mariner transposons are referred to as *TnSeq* methods. Tn5 transposons, on the other hand, do not target specific sequence motifs while exhibiting preference for GC-rich sites^4^. Tn5-based methods are called *TraDis* methods if the reads are sequenced directly after the PCR, or *HITS* if the PCR products are subjected to a size selection and a purification. The Tn5 is a common alternative as building Mariner-based library may be problematic for some species^1^. The downside is that the larger number of Tn5 insertions produce less saturated libraries. Another difference between the two transposons is the use of a restriction endonuclease in Mariner-based libraries. These enzymes, such as *Mme*I, cut a fixed-length sequence upstream from the site of insertion generating reads of equal length. The Tn5-based methods do not use this approach and produce fragments of various lengths potentially allowing for PCR bias^3^.

In the end each flavor of TIS experiments produces a collection of sequencing reads. Before the interpretation of TIS data can begin these reads need to be processed, mapped, filtered and converted into a form suitable for downstream analysis tools. TIS sequencing reads have complex structure as they include fragments of transposon backbone, primer annealing sites, molecular barcodes and other elements. For example, only ≈13 base pairs (bp) of a TnSeq read correspond the genomic region adjacent to the integration site—the portion that is mapped against the host genome—everything else needs to be stripped away before mapping. After read trimming and mapping resulting `BAM` datasets need to be filtered and converted into more compact representations such as, for example, `wig` format. Such *derived* datasets can then be paired with appropriate annotation datasets (representing the location of genes and functional elements across the host genome) and used as inputs to analysis tools.

A number of algorithmic approaches have been developed to facilitate TIS data analysis. These include Hidden Markov Model (HMM)-based methods for identification of essential sites as well as regression analyses utilizing gene saturation or runs of consecutive empty sites. These approaches are implemented in tools such as ESSENTIAL^5^, Tn-seq Explorer^6^, El-ARTIST^7^ suite, TRANSIT^8^, or Bio-Tradis^9^ (Table 1). The output of these tools—lists of genes classified as essential/non-essential or coordinates of regions enriched or void of insertions—need to be further processed by, for example, comparing results across conditions.

**Table 1.**
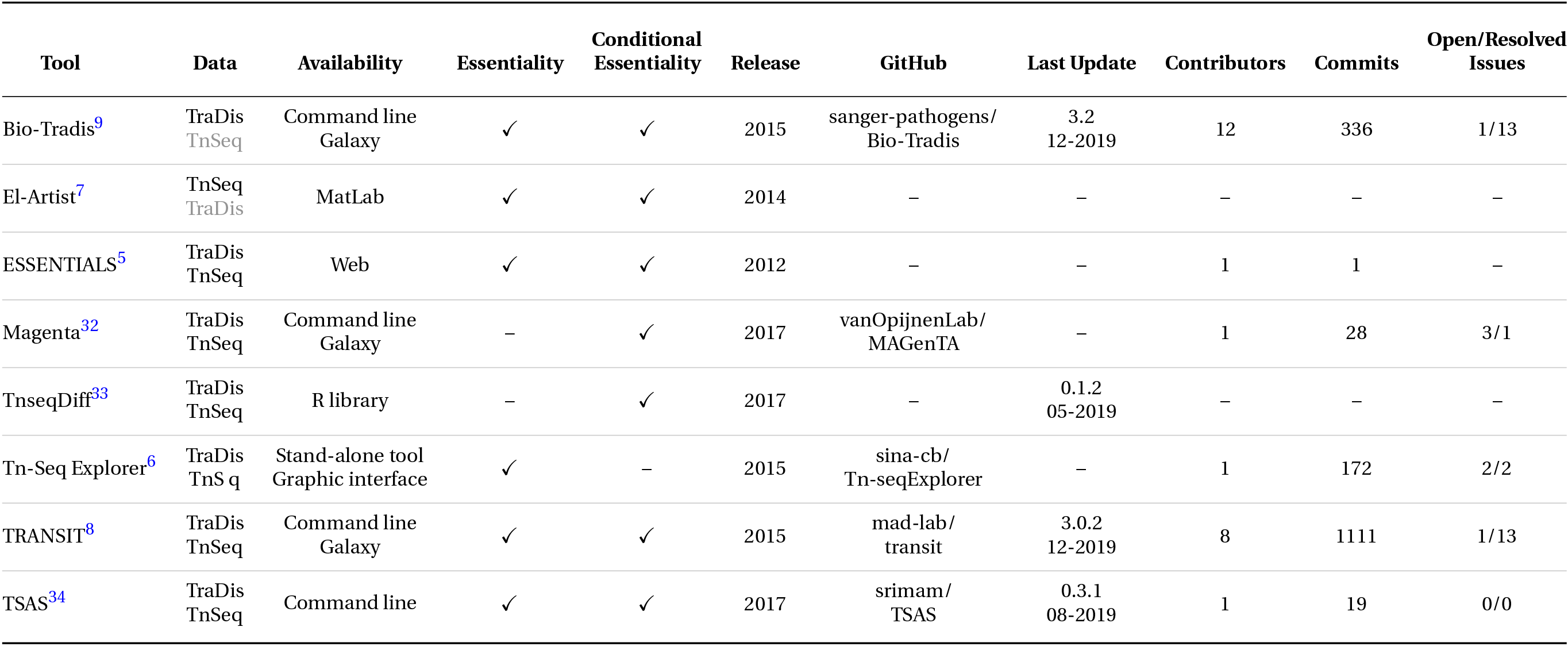
Tools availability and maintenance metrics : frequency of updates, number of contributors, commits, and issues. These metrics are indicators of the health of the tools : number of people maintaining the tools, frequency of update, responsivity of the developing team. Most of the tools can support both TnSeq and TraDis. The gray words indicate that the tools support the data in theory, but will necessitate to adapt the data. Across this list of tools, Bio-Tradis and TRANSIT seem well maintained. The other tool that have git hub repositories have only one contributor, which mean that the tool could stop being maintained if the one contributor change project. Among these 3, 2 have not been updated since the first release some years ago. Three tools have no Github directory, although ESSENTIAL has a svn repository. The svn logs shows a single commit in 2012.

The tools such as TRANSIT and Bio-Tradis offer powerful means for the analysis of TIS data. However, while they cover the essential step of data interpretation—it is just one part of a multi-step process involving, as we demonstrated above, trimming, mapping, filtering, and additional statistical analyses. In this manuscript we developed a set of comprehensive re-usable workflows for analysis of TIS data that include all analyses step from initial read processing to preparation of final figures for publications. To demonstrate the utility of our approach we reanalyzed data from two recent studies employing Tn5 and Mariner transposons. Add tools and workflows developed by us are publicly available to immediate re-use as described at https://bit.ly/gxy-tis.

## Results and Discussion

Our goal was to design a publicly accessible high throughput system for transparent and reproducible analysis of transposon insertion data. To devise and test our approach, we selected data from two recent studies: one performed in *Escherichia coli*^10^ and another conducted in *Staphylococcus aureus*^11^. The first study^10^ used the TraDIS approach^12^ based on a Tn5 transposon inserting with high-frequency into arbitrary sites within the target genome^13^. The second study^11^ used the TnSeq approach based on a phage-assisted Mariner-derived system that targets `TA`-sites within the host’s genome^2^.

### Analysis of TraDIS data

Goodall et al.^10^ used TraDIS to identify essential genes in *Escherichia coli* BW25113. TraDIS technique generates reads that contain experimental barcodes and segments of the transposon backbone in addition to the fragment of genomic DNA proximal to the insertion site. Before the reads can be used for mapping, required to identify locations of insertion sites, they need to be trimmed down to include only genomic DNA adjacent to the insertion site. In the case of the Goodall et al. study^10^ this has been done prior to submitting the sequencing data to the Short Read Archive (SRA - BioProject PRJEB24436), and thus the reads can be used directly for mapping without any preprocessing. We mapped the reads against the *E. coli* BW25113 genome (CP009273.1) and computed read coverage (see Methods) using the 5’-end of the reads as it is immediately adjacent to the insertion site^12^.

#### Regression on genes saturation indexes

To identify essential genes we proceeded to carefully re-implement the analysis performed by Goodall et al.^10^. These authors conducted a regression analysis on gene saturation indices by fitting known distributions to the distribution of insertion indexes. First, we computed gene saturation index *S*—a simple statistic calculated by dividing the number of insertions within a coding region (CDS) by its length. In this dataset, *S* is bimodally distributed with the first mode at low saturation and the second at high saturation (Fig. 1). This profile is coherent with the expected distribution of gene saturation in TIS studies with saturated libraries^1^. It is a mixture of two distinct distributions: one of essential genes and the other of non-essential genes.

**Figure 1.**
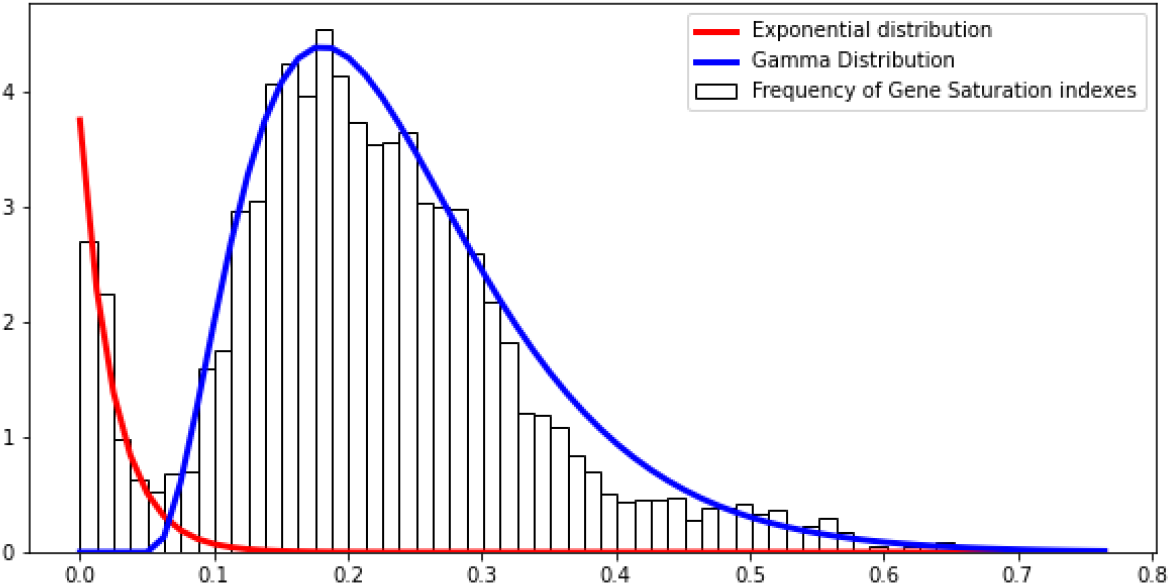
Regression analysis on gene saturation indexes for TraDIS dataset. The histogram represents the frequency of saturation indexes of genes. A gene saturation index is the ratio of insertion sites with insertions on the number of potential insertion sites. The regression uses the density of frequencies, as we are using density function to fit the data. A parametric regression algorithm is used to divide the bimodal distribution in two distribution. An exponential distribution is fit to essential genes, and a Gamma distribution to non-essential genes. Although the Gamma distribution seem loose on the higher saturation sides, the two distributions seem to fit the data nicely in the saturation range where the division happens.

To separate two distributions we performed a regression by fitting a bimodal distribution to our data. The bimodal distribution is composed of an exponential and a gamma distribution components corresponding to essential and non-essential genes, respectively. Using distribution parameters we then computed a probability of every gene being drawn from exponential or gamma distributions. A gene is said to be *essential* if the probability of it belonging to the essential distribution is *X* times higher than the probability of belonging to the non-essential distribution (and vice versa). *X* is a threshold that can vary between studies, and genes that do not meet this requirement are classified as “undetermined”. Goodall et al.^10^ used *X* = 12 as it was previously employed in another *E. coli* gene essentiality study by Phan et al.^14^. Using *X* = 12 we identified 364 essential genes (Table 1). We then compared our results with the list of essential genes identified by Goodall et al.^10^ as well as those from Keio (based on BW25113;^15^) and PEC (based on MG1655;^16^) databases (Fig. 2). It showed that several essential genes predicted by us and absent from the Goodall et al. are found on the other databases. To evaluate the effect of differences in threshold we compared results obtained with 4-fold (*X* = 4) and 12-fold differences (*X* = used above. There were little differences between the two thresholds. (Sup. Table 1). With *X* = 4 we missed eight essential genes and overpredict 24 genes. *X* = 12 misses 12 essential genes and overpredicts 23.

**Figure 2.**
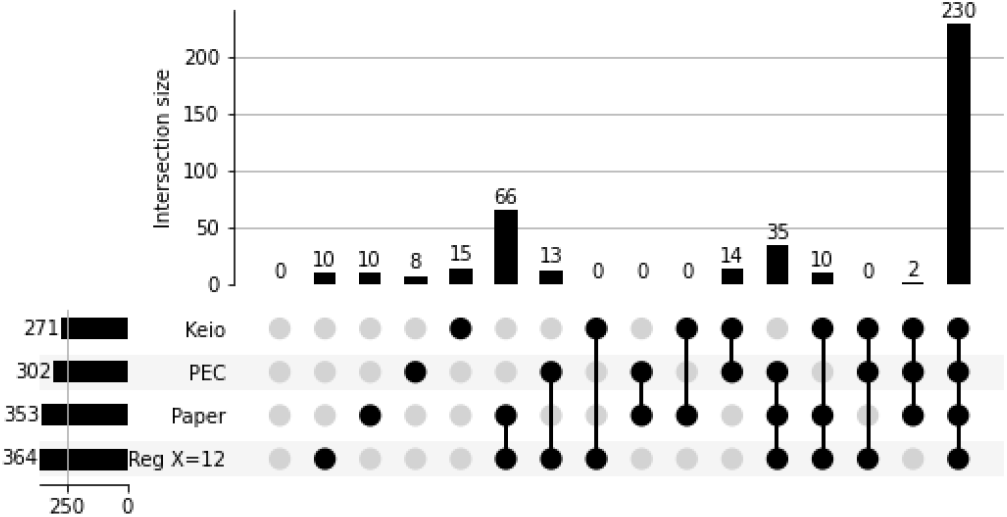
TraDIS - Upset plot comparing the regression results to Goodall et al. paper, Keio^15^ and PEC^16^ databases. We compared the list of essential genes resulting from our reproduction of the Goodall and al. study with their results and those identified by two external databases Keio and PEC. We can see that our replication predicts slightly more essential genes than the manually curated results provides, 364 genes against 353. Out of the 232 core essential genes identified in the study, we successfully identified 230. We identified most of the genes predicted by the paper that are absent in PEC or Keio (101 genes). Our replication identifies 13 essential genes identified in PEC but not in the original study in addition to the 35 genes identified everywhere but in Keio. The genes predicted by only one of the sources are of similar magnitude, a dozen genes each. They can be explained by manual annotation, parameter change, and domain essential genes.

#### Automatic Regression with Bio-TraDis

Next, instead of performing manual fitting we employed Bio-Tradis^9^ toolkit. It identified 398 essential genes, and classified the rest as “ambiguous”. We compared the essentiality prediction of Bio-Tradis with the previous regression analysis (Fig. 3C). Among the 353 essential genes listed by^10^, 351 were also identified by Bio-Tradis. It predicted 47 additional essential genes, among which 21 are also identified by the hand-fit regression. The results were very similar for gene essentiality, whether we used the automated or hand-fit method.

**Figure 3.**
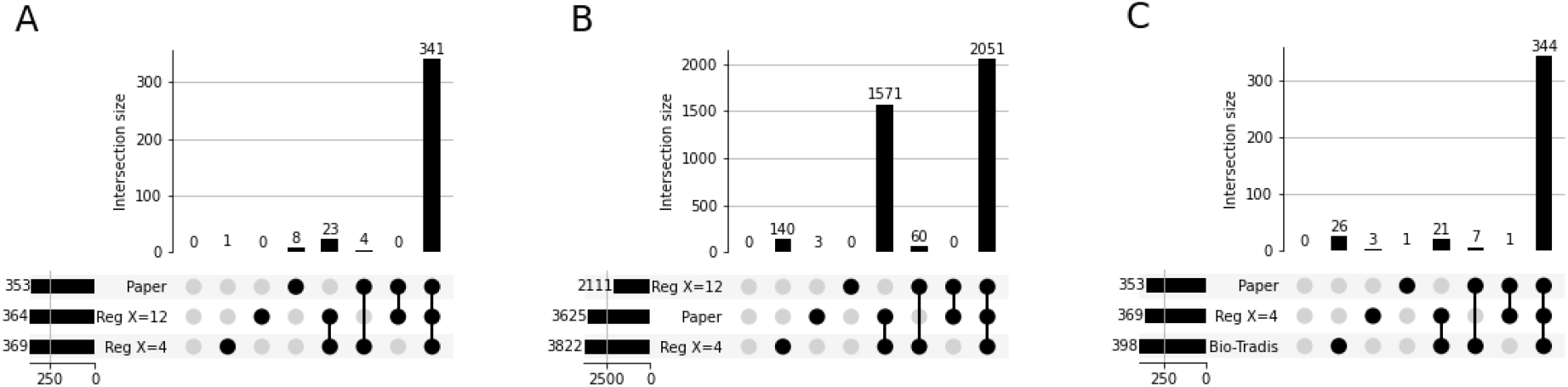
TraDIS - Comparison of methods for genes classification by regression analysis. A/ UpSet plot comparing the essential genes predicted by each threshold with the curated list. There is little difference between the two thresholds for essential genes prediction. 341 essential genes are identified in the three sets. Using a threshold of 4 predicts five additional genes compared to a threshold of 12, four of which are also identified in the curated list. B/ UpSet plot comparing the non-essential genes predicted by each threshold with the curated list. When looking at the non-essential gene list, we see that using a threshold of 12 misses 1570 non-essential genes identified in the paper and with a regression using a threshold of 4. The latest predicts 140 non-essential genes that are unclassified in the two other sets. C / UpSet plot comparing the non-essential genes predicted by using hand-fit curves, automated fit with Bio-TraDis, and the curated list. The two regression methods identify 344 of the essential genes predicted by the publication. One gene is missed by the two methods, and eight are identified by only one of the regression analyses.

#### Classification based on rows of empty sites

In addition to a regression performed to identify essential genes, Goodall et al. analyzed the density of transposon insertion in the genome and describe the probability of observing rows of empty (void of transposon insertion) sites^10^. This metric can be used to detect essential genes as well, and such a method is implemented in the Transit suite^8^ with the *Tn5Gaps* tool. Comparison of *Tn5Gaps* results to the Goodall et al. predictions (Fig. 4) showed that Transit predicted 124 additional essential genes, and 331 predicted essential genes were shared with the published results. The list of predicted non-essential genes generated by Transit, on the other end, was very close to the Goodall et al. results.

**Figure 4.**
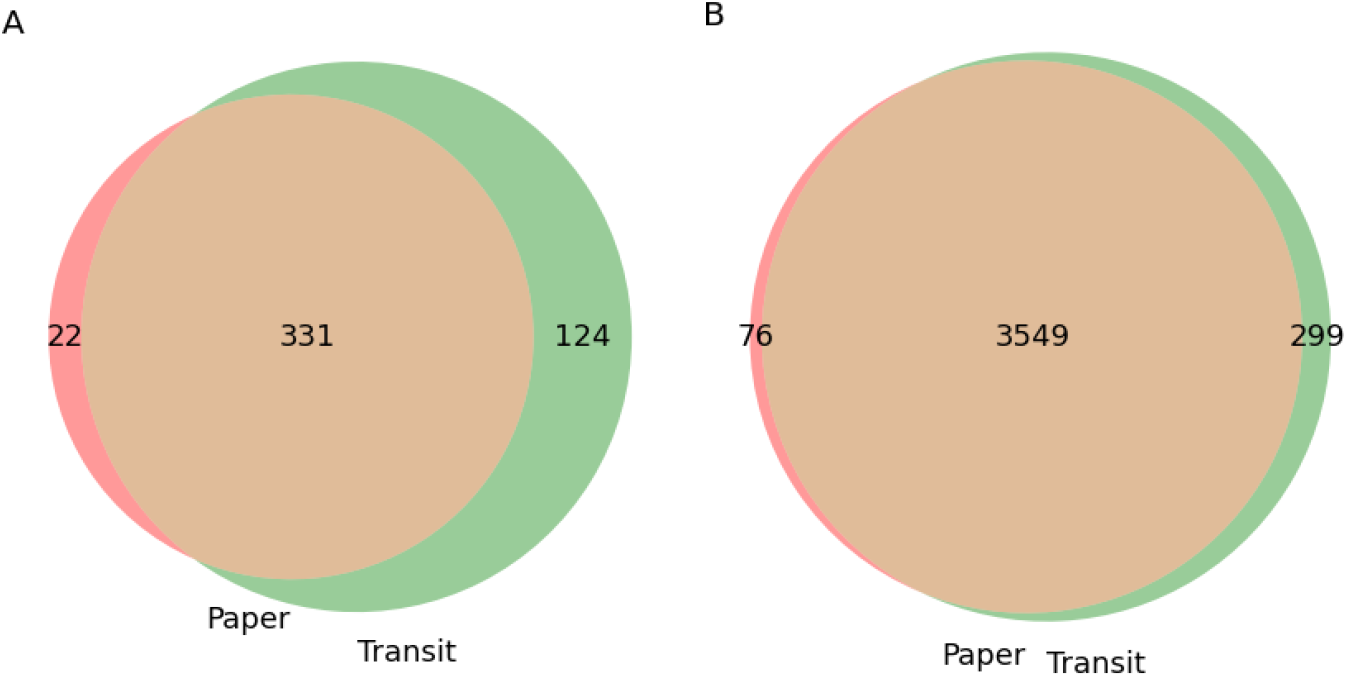
TraDIS - Comparison of genes predicted by the Gumbel tool in Transit and those published in the paper. A/ Essential genes. Transit seems to largely over-predict essential genes. 124 genes are identified as essential in addition to the 331 genes also identified as essential by manual curation. B/ Non-essential genes. Transit prediction of non-essential genes overlap well the genes predicted as non-essential in the paper.

#### Comparison of the regression and Tn5Gaps results

Finally, we compared the results of the regression, using the two thresholds, and the Tn5Gaps method implemented in Transit (Fig. 5). We could observe that Transit overpredicted essential genes (Fig. 5 A). Among the 124 genes identified by Transit but not by the Goodall et al study, 108 were identified by Transit alone. Some of the overpredicted genes were domain essential genes (only part of the gene is free of insertions), or smaller genes overlapping an empty region. Most of them, however, did not seem to contain essential regions but showed sparse insertions. When looking at non-essential genes predictions, both methods were very close to the paper (Fig. 5 B). The regression predictions were overall closer to the manually curated results than Transit regardless of the chosen threshold.

**Figure 5.**
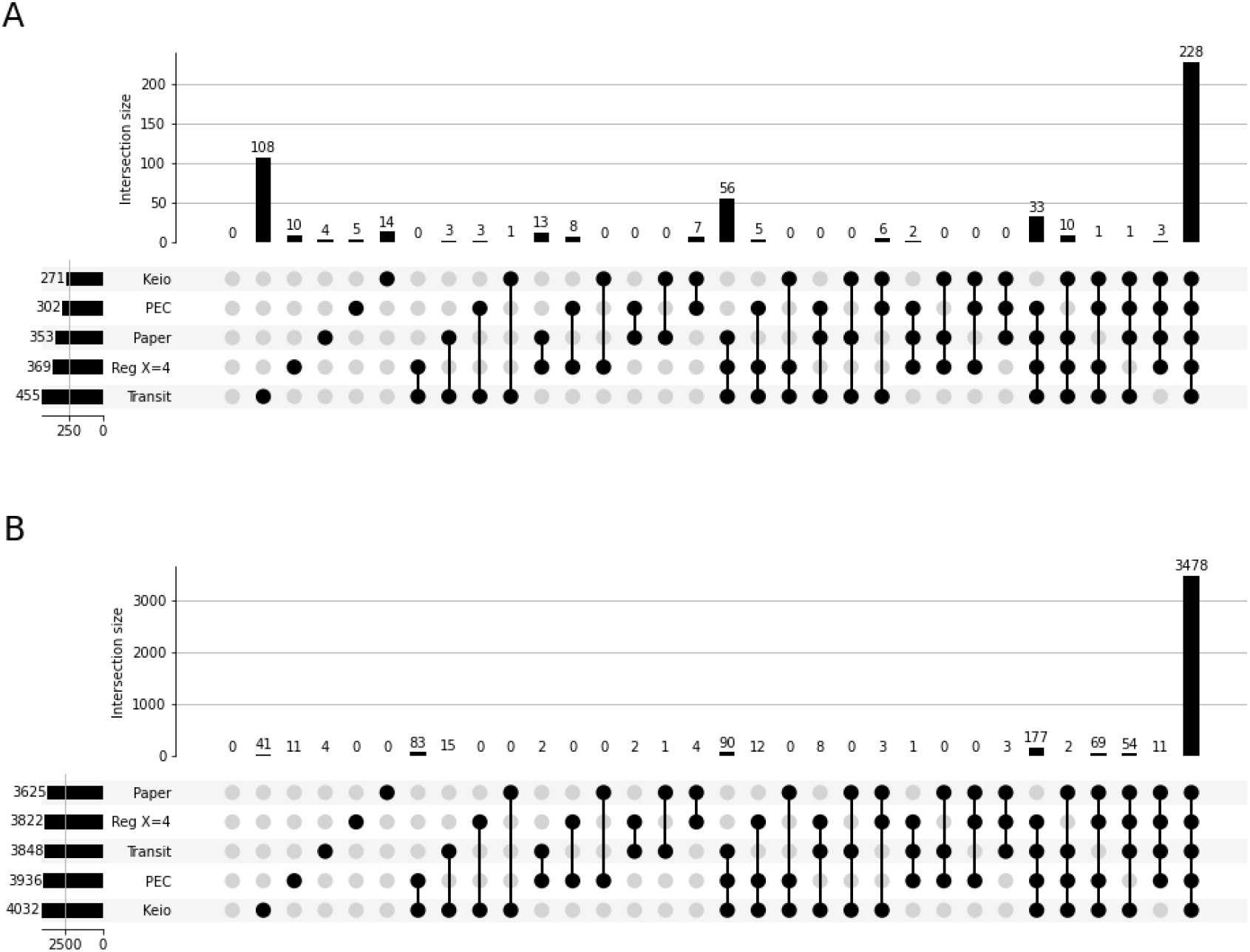
TraDis - Comparison of regression results, Transit results, external databases, and the curated list of essential genes. A/ Comparison of essential genes predictions. We can see that Transit over predict the number of essential genes compared to databases as well, 108 are classified as such that does not appear in any other source. B/ Comparison of non-essential genes predictions. All sources have similar results for non-essential genes identification. 3478 genes are always identified. The second-largest set of genes are genes identified as unclassified by the manual curation only. These could correspond to domain-essential genes or genes that impact growth.

#### Read count normalization

The transposon insertion sequencing datasets can be normalized for three factors: (i) positional read bias,(ii) differences in sequencing depth, (iii) stochastic differences in library diversity^1^. To evaluate the impact of normalization on the essentiality prediction, we compared the results obtained without normalization and using a normalization method provided in the Transit toolset. In particular, we used the “TTR” method recommended by the authors, to remove outliers and normalize for differences in saturation^17^. For both analyses and both essential and non-essential genes, the results were identical regardless of the normalization.

### Analysis of TnSeq data

We applied the analysis approach developed on TraDis data to a Mariner-based TnSeq dataset produced by Santiago et al.^11^. The methodology pioneered by these authors allows transposon-mediated insertion of promoters into the target genome^2^. As a result this technique provides two types of readout. First, similarly to TraDis, lack of TnSeq reads mapping to a genomic locus indicates its functional importance. This readout—lack of reads—can be used for identification of essential genes. Second, promoters contained within insertion constructs may affect neighboring genes. Because promoters act in a directional fashion sequencing reads derived from these insertions exhibit strand bias. The second type of readout—regions where the majority of reads map to one of the two strands—allows finding genes those expression change is beneficial given an experimental condition. In this study we focused on the first type of TnSeq readout to identify essential genes. The data produced by Santiago et al.^11^ contain raw reads (SRA BioProject PRJNA417822) for 82 experimental conditions with varying number of replicates. We used the control condition (containing 14 replicates) to develop reproducible strategies for read preprocessing and control for noise in the data.

#### TnSeq data preprocessing

Sequencing reads produced by Santiago et al.^11^ contain transposon backbone and auxiliary sequences that need to be removed prior to analysis. In addition, the reads contain molecular barcodes that identify constructs containing different promoters. The genomic portion of the reads, that is ultimately mapped against the host genome, is only 16-17 bp in length because TnSeq protocol uses *Mme*I restriction endonuclease. *Mme*I cleaves DNA 18 nucleotides downstream of recognition site. After mapping against the *Staphylococcus aureus* (CP000253.1) we computed coverage at 3’-end of the reads only as this corresponds to the position of the insertion site (Fig. 6 B). Finally, we compared the coverage information with the position of all `TA` dinucleotides found in the the *Staphylococcus aureus* genome.

**Figure 6.**
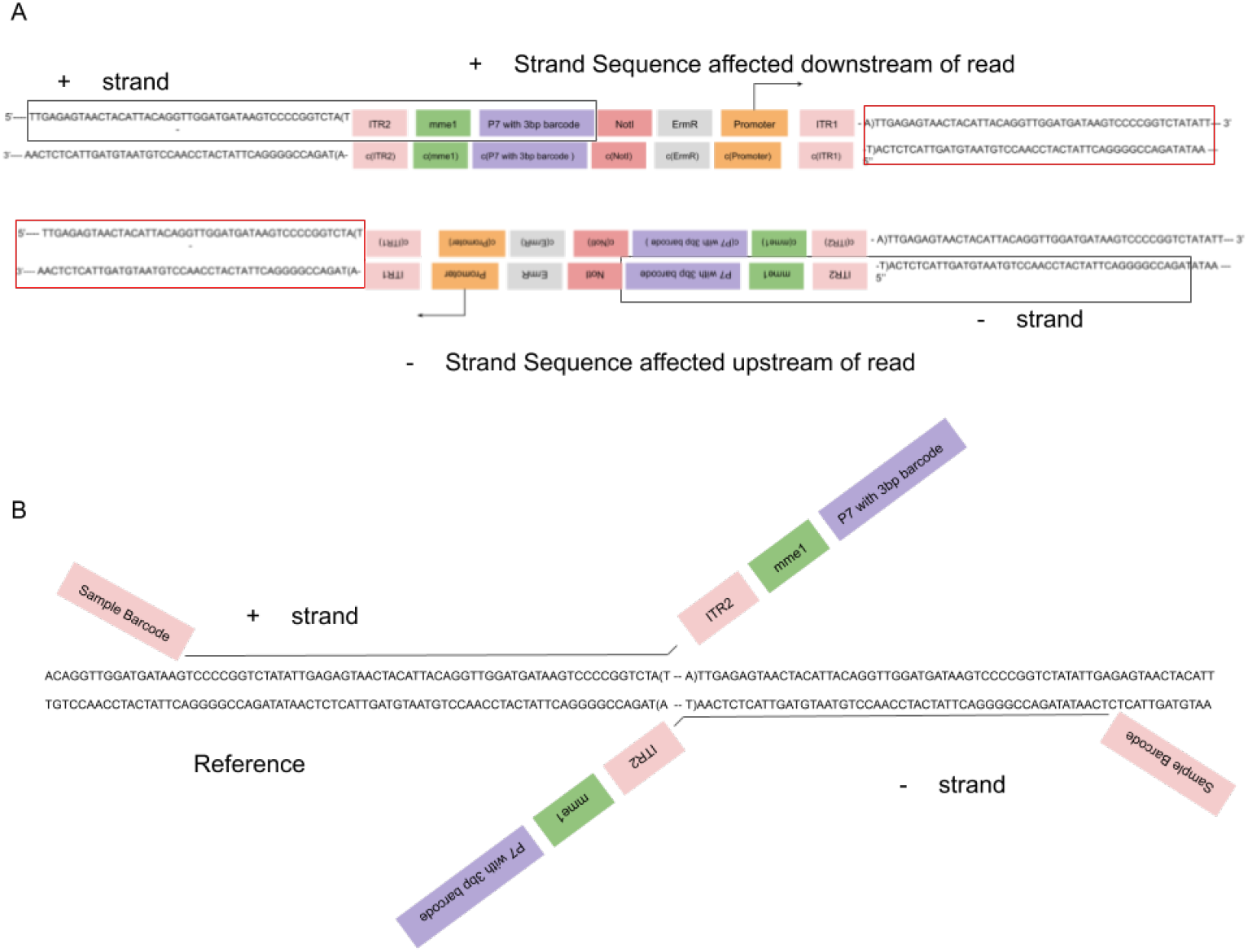
TnSeq - Structure of transposon insertions and reads in Santiago et al.^2^ technique. A/ Structure of a transposon construct insertion. The transposon can insert in two directions since a TA site is identical in both strands. The MmeI insertion in the transposon allows reads of uniform size by cutting DNA 20 bases upstream of the transposon. Most of the inserted promoters are pointing toward the downstream regions. B/ Structure of the reads. In addition to the fragment of 20bp upstream of the MmeI site, the reads contain an Illumina adapter P7 containing a 3bp barcode specific to the type of construct, and an Illumina adapter P5 containing a sample barcode at the 5’ end. The read also contains the MmeI transcription site included in the ITR region. We can see from that structure that the position of the insertion site is aligned with the 3’ end of the read.

#### Regression on the TnSeq dataset

The regression is the method that gave us the best results with TraDis data. However, TnSeq data are distribution of the saturation index *S* is different for TnSeq data (Fig. 7). Here we used exponential distribution for the essential genes and normal distribution for the non-essential genes. When correcting the method to use appropriate distribution, we could compare the efficiency of the different thresholds (Fig. 8 A). The threshold choice had more profound effect of TnSeq data: the use of *X* = 4 as a threshold lead to the prediction of 480 genes, which was higher than the number of essential genes expected, and more than predicted with the use of a threshold of *X* = 12 (Sup. Table 2). When compared with the results of the publication^2^, most of the essential genes and half of the domain essential genes were identified by the regression analyses (Fig. 8). The use of a threshold of *X* = 12 produced more false negative results than *X* = 4, and as many false positives. The prediction of non-essential genes was very close to the published results when using either threshold (Fig. 8 B). Overall, the choice of the threshold in this analysis appeared to change the essentiality prediction significantly. A threshold of 12 appeared too stringent in that case, perhaps due to looser regression curves compared to TraDis data.

**Figure 7.**
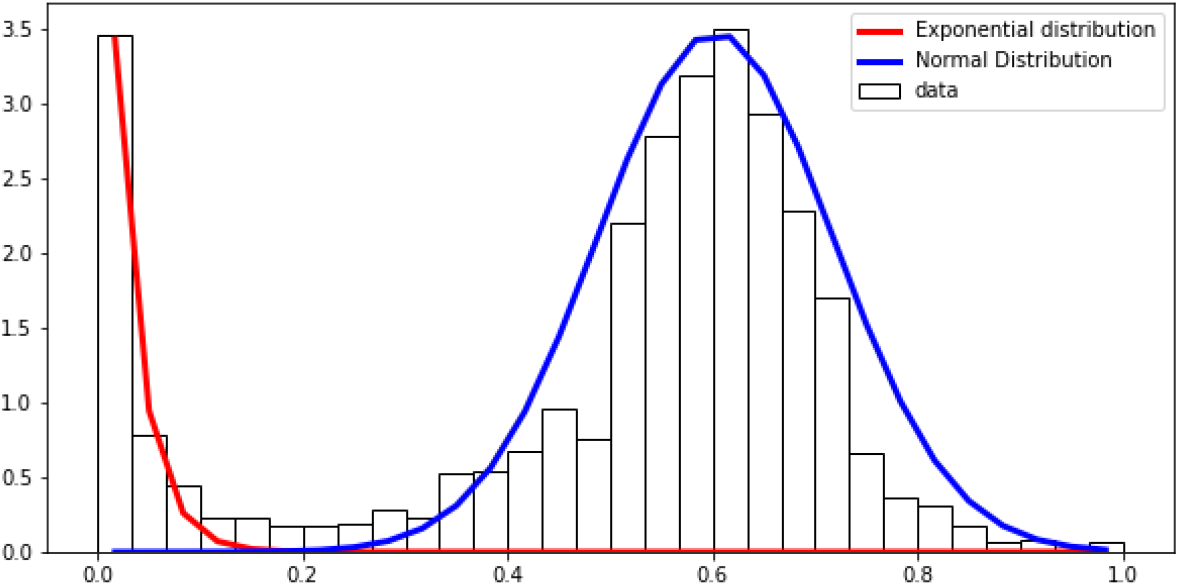
TnSeq - Regression on the frequency of gene saturation indexes for HiMar transposon and S. aureus. The Exponential distribution and the Normal distribution seem to fit the general shape of the distribution, despite some lose fit around the quartiles of the normal distribution. There is no intersection between the two distribution; however, some genes have a null probability of belonging to either population.

**Figure 8.**
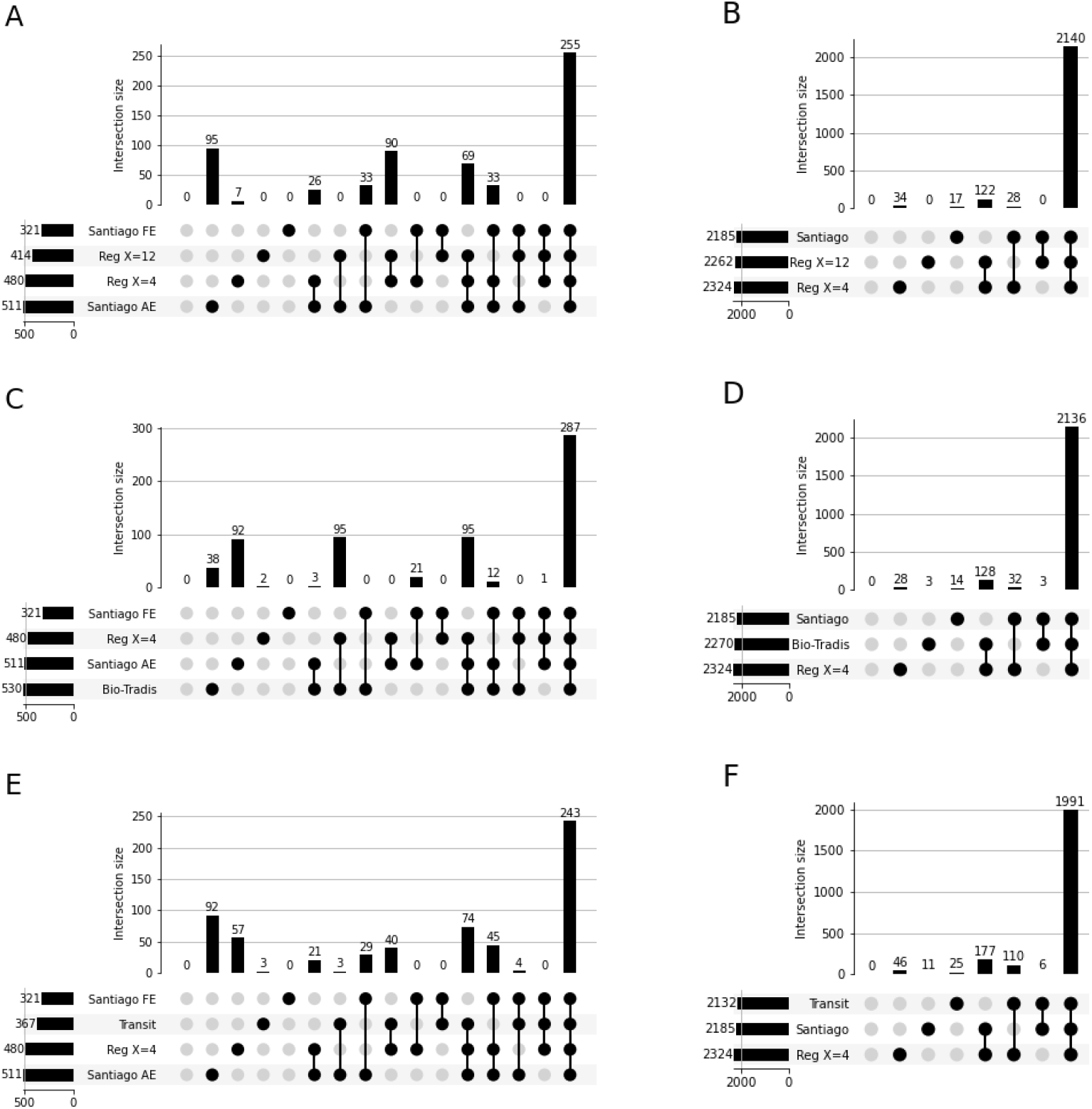
TnSeq - Comparison of the essential genes prediction per method. Santiago FE refers to the genes that are identified as fully essential in the Santiago et Al. paper. Santiago AE refers to both fully essential and domain-essential genes. The classification of the regression results has been performed with thresholds of 4 and 12. A/ Comparison of regression on gene saturation index with the genes identified as essential in Santiago et al.^2^. 255 genes are shared between the regressions and the paper full essential genes list. B/ Comparison of regression on gene saturation index with the genes identified as non-essential in Santiago et al. The methods do not differ much on non-essential genes predictions, 2071 genes are identified by the paper and the regression, regardless of the chosen threshold. C/ Comparison of Bio-Tradis prediction of essential genes with the hand-fit regression and paper predictions. 287 genes are common to the three methods, and Bio-Tradis overpredicts 32 genes not identified by other methods. D/ Comparison of Bio-Tradis prediction of non-essential genes with the hand-fit regression and paper predictions. The results are very similar. 197 genes are predicted by both regressions that are not classified as non-essential by the study. E/ Comparison of Transit results, regression results, and genes identified as essential in Santiago et al. We are using the results of the hand-fit regression using a threshold of 4, to be more relaxed, as the threshold of 12 provides more stringent results and Bio-Tradis overpredicts essential genes. F/ Comparison of Transit results, regression results, and genes identified as non-essential in Santiago et al. The three methods are in accordance with the non-essential genes predictions, 1923 are common across all results.

#### Automatic Regression with Bio-TraDis on the TnSeq dataset

We compared the results of our regression analysis to the automated regression performed with the Bio-Tradis toolkit (Fig. 8 C and D). Bio-Tradis predicts a larger number of essential genes compared to in Santiago et al.^2^. Our method had a better performance than the tool for the prediction of essential genes. The tool had a slightly better rate of true-positive compared to the hand-fit regression, but a worse rate of false-positive. When we looked at the results of non-essential genes predictions, the two methods performed equally.

#### Classification of genes based on rows of empty sites on the TnSeq dataset

Using the same method as Tn5gaps, the Gumbel tool in the TRANSIT suite is designed for gene essentiality prediction based on Mariner transposons. As with the regression method, we did not differentiate essential or domain essential genes. Therefore, we compared our results both with the essential genes, and essential and domain essential genes identified by the essentiality study with TnSeq data^2^ (Fig. 8 C). Transit and the regression analysis identified half of the full essential genes, about a third of domain-essential genes. Overall, transit performs more poorly that the other methods. It could be due to the higher sensitivity of Transit to gene sizes compared to saturation indexes. The minimum size of essential regions needed for the identification of essential genes by Transit needs to be shorter than a gene to be found significant. Many of the domain essential genes were not identified by any method. The identification of domain essential genes by Transit or the regression method is dependent on the size of the essential domain. Too small, and it would not appear as significant for Transit, and would not impact the saturation index enough to be classified into the essential category.

When looking at the non-essential genes, we could see that all the methods provided very similar results (Fig. 8 D). It seemed to indicate that only the essential genes predictions were impacted by this variation in condition. Some genes may have been going from growth defect to entirely essential depending on the conditions. It would have impacted their classification into essential genes, going from low insertion rate to no insertions, without them having sufficient saturation index to be classified as non-essentials.

#### Prediction of Essential genes using HMM on TnSeq Data

TnSeq analyses often use a third method in addition to the one used in the TraDis. Hidden Markov Model (HMM) methods consider each potential insertion site independently of the previous states to predict the state of the next position and attribute costs of changing from one state to another. This method allows predicting regions with the same state. It is a standard method available in several tool suits, like EL-ARTIST, TRANSIT, ESSENTIALS, Tn-seq explorer, and MAGENTA (See methods). Our goal was to identify a tool that facilitate reproducibility and transparency of analyses. For these reason we chose Transit suite to perform an HMM analysis of the TnSeq Data—it has a permissible open source license and is actively maintained (Table 1). This tool uses HMM to classify each site into four states: Essential, Non-Essential, Growth Defect, and Growth Advantage. This method returned about two thousand essential genes, which is more than the expected 350 to 400 genes. This result could be explained by the fact that Santiago data may be too sparse for the HMM, which is sensitive to zero-inflated datasets.

#### Comparison of the different methods for essentiality prediction on TnSeq data

We compared the results of the regression, using a threshold of 4, and the Gumbel with the original study, and other similar essentiality studies (Fig. 9). We did not include the HMM analysis, as the results were inconclusive. We compared our predictions to two other published TnSeq studies: Chaudhuri et al.^18^, and Valentino et al.^19^ (Fig. 9). We could observe a core of 229 genes reported as essential across all studies. Overall the results provided by Transit and the regression analysis were coherent with the results expected from essentiality analyses compared to other studies performed in different conditions. Transit prediction was closer to the other studies, with only two reported essential genes not predicted by other methods. The regression method identified 54 additional essential genes compared to other studies. The additional 89 predicted in the Santiago analysis could be explained by its distinction between essential and domain essential genes. For this TnSeq dataset, Transit performed better than the other methods.

**Figure 9.**
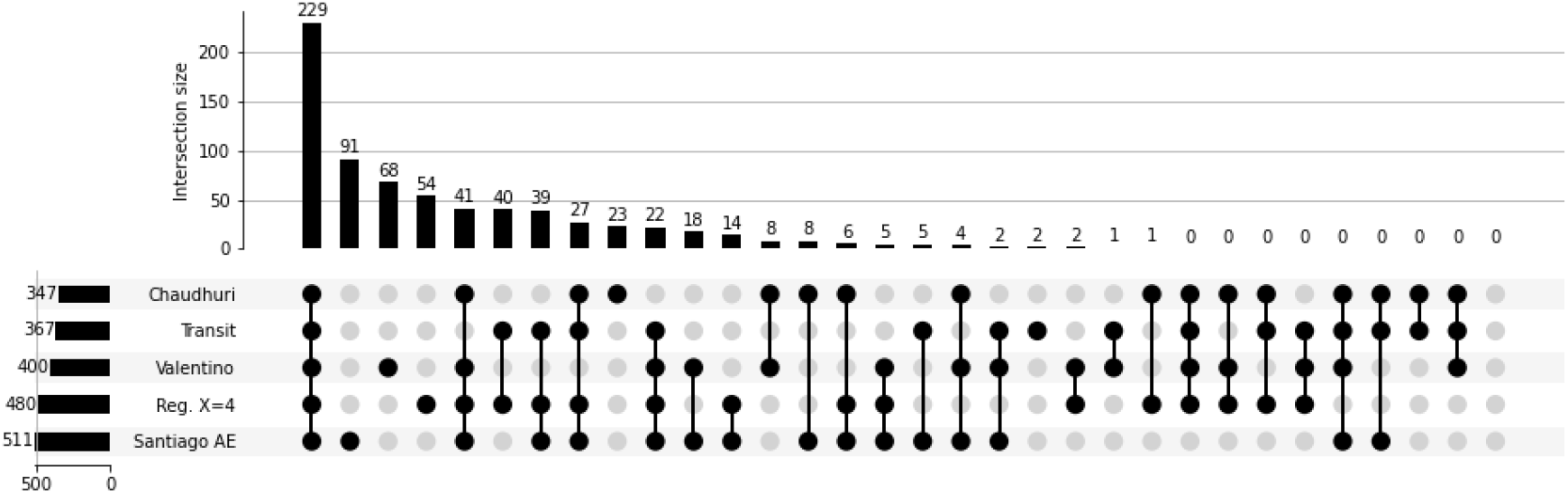
TnSeq - upSet plot of the essential genes predicted by the different methods, the genes published by the reference paper^2^, and other studies of gene essentiality in S. aureus. 229 genes are identified as a “core” of essential genes between studies. As Santiago et al. paper differentiates domain essential and full essential genes, it shows the largest difference compared to the other studies with 89 genes that are not found in other sets of essential genes. Transit is the closest to all other studies, with only two extra genes predicted, against 54 predicted by the regression analysis.

### Implementation in Galaxy

We implemented the analysis workflows in Galaxy^20^, including preprocessing, alignment, read counts, and essentiality prediction. We also developed and deployed a comprehensive tutorial explaining application of our system to TIS data^21^. Despite being very similar, the two types of data, TnSeq and TraDis, require distinct workflows, due to the difference in read structure and coverage analysis. We constructed these workflows and made them available in the Galaxy main instance (usegalaxy.org; Supplemental Figs. 1 and 2). The TraDis workflow includes two parallel tracks (Supplemental Fig. 1). One track performs essentiality analysis with Bio-TraDis^9^ and the other one with TRANSIT^8^. The latest include the mapping of reads with Bowtie2^12^ and the analysis of coverage with bamCoverage tool^22^. The TnSeq Workflow is more complex than the TraDis one, in part due to the read prepossessing needed with this dataset. Finally, steps have been included to replace the tool *Reads to counts* from the Bio-Tradis suite, which is not suitable for TnSeq data. This workflow uses Bowtie to map reads, as it is more adapted to very short reads than Bowtie2. Data are imported into Galaxy using the SRA tool fasterq-dump^23^. All the steps are highly adaptable to different read structures and sequencing technology agnostic. The entire workflow runs in approximately half a day on TnSeq data from^2^. The most time consuming steps were splitting by barcodes and alignment. The availability of Jupyter in Galaxy permitted the in-depth analysis of the final data. We included these notebooks in galaxy histories containing this paper analyses (See Supplemental Information). We used them to perform regression analyses and to compare the results of different methods with other published studies.

## Conclusions

The goal of this study was to develop end-to-end workflows for the analysis of transposon insertion sequencing data. TIS studies are used to identify genes essential to bacterial growth in specific conditions or to detect genes causing growth defect or advantage. While the experimental aspects of the technique are constantly evolving, there is no consensus on the way to perform the essentiality data analysis as a number of different approaches is described in the literature. These can be broadly classified into three categories. The first group of methods is based on a regression analysis of the gene saturation indices. The second type of analysis uses runs of consecutive sites with no transposon insertion. Finally, the third group consists of HMM-based methods. We added tools representing each of these categories to Galaxy toolkit.

To test performance of these methods we selected two different datasets. The first one is a TraDIS dataset utilizing Tn5 transposons that can potentially insert at any genomic position. The second dataset used Mariner transposons inserting at `TA` sites. The choice of transposon impacts the overall library saturations: TraDis data is more sparse than TnSeq data because Tn5 transposons have many more potential insertion sites compared with Mariner transposons. The TnSeq library has been built using different types of promoter-containing construct. We performed the analyses using either the control constructs, that do not contain any promoter, or all constructs as replicates. The second solution provided more reliable results by increasing library saturation.

Regression analysis produced results that were in good agreement with published data as well as with databases of essential genes. We performed the manual regression in a Jupyter notebook in Galaxy. We also used the Bio-Tradis^9^ tools suite to perform an automatic regression. The manual method adds the burden of the choice of distribution on the user. On the other hand, it provides more flexibility as differently saturated libraries may present different profiles that would change the distributions. When the appropriate parameters are determined, the manual method was highly reliable on datasets we tested.

HMM analyses produced inconclusive results in our hands. In both cases, the data were too sparse for HMM to identify stable regions (consecutive insertion sites with the same state that cover a genomic region large enough to be considered significant biologically). These results were expected for the TraDIS dataset, due to the sparse nature of the libraries, but were more surprising for the TnSeq data.

The workflows implemented in Galaxy are using robust open-source tools. The tools used explicitly for TIS analysis are open source and well maintained, ensuring reproducibility and transparency of analyses. The workflows include Jupyter notebooks for exploratory analyses without losing the history tracking in Galaxy that allows traceability and sharing. The combination of open source tools and Jupyter notebooks provides a complete and flexible workflow that can be easily modified to fit any analysis need.

## Methods

### Selecting appropriate tools

First we assessed the status of existing tools for the analysis of TIS data (Table 1). This information is essential for tool selection as it identifies actively maintained tools that will be supported in the future. Based on this analysis only Bio-Tradis, Magenta, and Transit are actively developed, maintained, and regularly released (Table 1).

### Alignment of TraDis data

The reads have been trimmed to remove low-quality (Phred score < 20) bases at the end of the reads with Trimmomatic^24^. The reads were mapped using Bowtie2^12^ against the reference genome of *Escherichia coli* BW25113 (CP009273.1). The coverage of the genome was computed with BamCoverage, from the Deeptools suite^22^. In our case, we were interested in identifying only insertion points. For that reason, we computed coverage using 5’-ends of the reads only.

### Transit for TraDis data

Tn5Gaps is the Gumbel tool adapted for TraDis data. These methods perform a gene by gene analysis of essentiality based on the longest consecutive sequence of potential sites without insertions in a gene. This metric allows identifying essential domains regardless of insertion at other locations of the gene. A We ran Transit with default parameters, but changed the normalization method and selected not to normalize the counts. Transit offers two ways to deal with replicates: either the counts of all replicates at each site are summed, or they are averaged. Using the sum of counts provides a better saturation of the library: a site that might not have been impacted during the initial transfection might have been impacted in another sample. In the case of TraDis data, where the saturation tends to be low, due to a large number of potential sites, the use of the sum of counts provides a better resolution.By averaging the counts at each site, we significantly decreased the noise.

### Saturation Indexes

Saturation is defined by the ratio of the number of sites impacted by insertions on the total number of sites able to receive an insertion in the gene. If the library is sufficiently saturated, we should observe two distinct distributions. The essential genes distribution has a low average saturation, and the distribution of the non-essential genes has a higher average saturation.

### Regression on TraDis data

Regression is a statistical analysis aiming to estimate the relationship between variables. It provides a function modeling this relationship. In our particular case, we are trying to identify the models behind the observed distribution of gene saturations. Formalizing these models then allows calculating the probability of each gene to belong to either one. We performed this regression using the python library scipy^25^.We started by defining the probability density function (*pdf*) of the gene saturation as the sum of two known *pdf* s: here, an exponential and a *γ* distribution. We anchor the regression by providing expected parameters. Goodall et al.^10^ did not provide the parameters and thus we had to estimate them. We attempted plotting several distributions and correcting the parameters to make them look like our data as much as we could (See Jupyter notebook). Once we approximated the parameter, the fitting function fits the data and returns the corrected parameters and their standard deviations for the two distributions.

### Classification of TraDis data

Once we identified the two distributions, we calculate the probability of genes to belong to each of them. This probability is calculated using the probability density function of each class (library *scipy* in python). *pdf* s are function whose area under the curve in the interval *X* = [*a*, *b*] is the probability of a random value of the distribution to belong to the interval [*a*, *b*]. When used for a single value, it also provides a relative likelihood that the value *n* belongs to the distribution (the absolute likelihood of *n* is null, since our variable is continuous).

To decide between the two categories, we select the most likely category if the difference between the two probabilities is significant. Formula (1) defines the significance thresholds for gene classification :

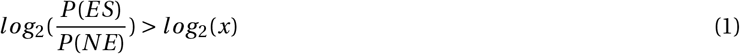

The threshold is calculated as the *log*_2_(*x*), where *x* is the number of times the likelihood for a mode must be superior to the likelihood of the other one. The use of a logarithm allows ignoring the direction of the ratio for the decision calculation. Genes whose difference had not been judged significant are classified as undetermined.

### Bio-TraDis on TraDis data

The Bio-Tradis toolkit provides a comprehensive set of tools for preprocessing reads from TIS. Our data are already exempt from transposon sequences, and we used the dataset provided as input for a read mapping step, followed by the count of insertion per gene, and gene essentiality predictions. The mapping is done using bwa, with a minimum mapping quality of 0 and default values for the other parameters. The toolkit does not handle replicates, so we merged the datasets after the mapping by adding read counts at every position.

### Pre-processing of TnSeq data

The reads published still contain transposon sequences that need trimming. The transposon sequence includes not only primers but also barcode sequences that separate the constructs containing different promoters. We use Cutadapt^26^ to separate the reads of each construct. We then trimmed the transposon sequence downstream of the construct barcode, the sequence including the ITR and the MmeI site, the sample barcode at the beginning of the reads, and finally, the low-quality bases before alignment and calculation of coverage.

### Alignment of TnSeq data

The reads have been aligned to the reference genome of *Staphylococcus aureus subsp. aureus NCTC 8325* (CP_000253.1) using Bowtie. Bowtie^27^ has been selected instead of Bowtie2^28^ for this dataset, as it is recommended for very short reads (shorter than 50 bp). We enforce alignment with no mismatches. It is necessary as we are working with very short reads of 16-17 bp, which is the minimum length for the majority of reads to have a unique alignment^29^.

### Counts for TnSeq Data

The coverage of the genome is computed with BamCoverage, from the Deeptools suite^22^. The coverage is calculated with 310 an offset of −1, meaning that the read is counted only at the 3’ position, at the TA site position (Figure 7B). In this study, we have seven datasets for each biological replicates: one dataset corresponding to control constructs, and six corresponding to transposon containing a promoter. Contrary to the TraDis data, where the Tn5 transposon inserts everywhere, Mariner transposons insert only at TA sites. To compute the gene saturation and row of empty sites, we need to use the coverage of all TA sites, whether it is nul or not. It is accomplished by merging the coverage files with the positions of TA sites calculated with the Nucleotide subsequence search tool available in Galaxy^30^. The resulting file is a tabular file with a column containing the position of the read (leftmost site), and another containing the counts of reads aligning at this TA site.

### Bio-Tradis TnSeq data

The Bio-Tradis toolkit provides a comprehensive set of tools for preprocessing reads from TIS. This preprocessing requires to provide the sequence of the transposon located at the beginning of the read. Sequences needing trimming in our reads are located on both sides of the reads and are variable between constructs. Bio-Tradis includes a script for read alignment that provides both the mapped reads and the insertion counts at each nucleotide. The scripts doe not provide the option to choose which end of the read should be used to attribute the insertion at a nucleotide. To circumvent this problem, we used the counts generated by our workflow as input for Bio-Tradis essentiality analysis.

### Transit for TnSeq data

We ran HMM and Gumbel tools on the TnSeq data. The HMM results were inconclusive, probably due to the low saturation of each sample after we divided the data based on transposon constructs. We ran the Gumbel method with default data, except for the option to not normalize data and the choice of replicates handling method. For this dataset, we are using the mean instead of the sum because of the large number of samples. While using the sum increases the library saturation, it is more sensitive to artifacts than the mean. If one read align by mistake at a position in 30% of the 84 samples, the mean would be 0 when the sum would be 15. it makes a big difference in the metrics used for essentiality prediction.

### Regression on TnSeq data

The regression on TnSeq data follows the same protocol as the one used for TraDis data. The difference is that we did not have any information about the type of distribution we expected. We added a step of distribution selection to the parameter estimation. We used a book describing the different distribution to select those who appeared close to the shape of our data^31^. We first started with the distribution of non-essential genes (Figure 7 A). The distribution shows no skew, and seem to correspond to a normal distribution. An exponential distribution seemed to be the most appropriate to describe essential gene saturations.

## Acknowledgments

Usegalaxy.org efforts are funded by NHGRI Grant U41 HG006620, NSF ABI Grant 1661497, NIAID grant R01 AI134384. We would like to thank the entire Galaxy team for supporting this effort.

## Supplemental data

**Supplemental Table 1.** Gene essentiality prediction of regression for both thresholds, Tn5 with E.coli.

| Threshold | $\log_2(4)$ | $\log_2(12)$ |
| --- | --- | --- |
| Essential | 369 | 364 |
| Non-Essential | 3822 | 2111 |
| Undetermined | 112 | 1828 |

**Supplemental Table 2.** Gene essentiality prediction of regression for both thresholds, Himar1 and S.aureus.

| Threshold | $\log_2(4)$ | $\log_2(12)$ |
| --- | --- | --- |
| Essential | 480 | 414 |
| Non-Essential | 2324 | 2262 |
| Undetermined | 37 | 165 |

**Supplemental Figure 1.**
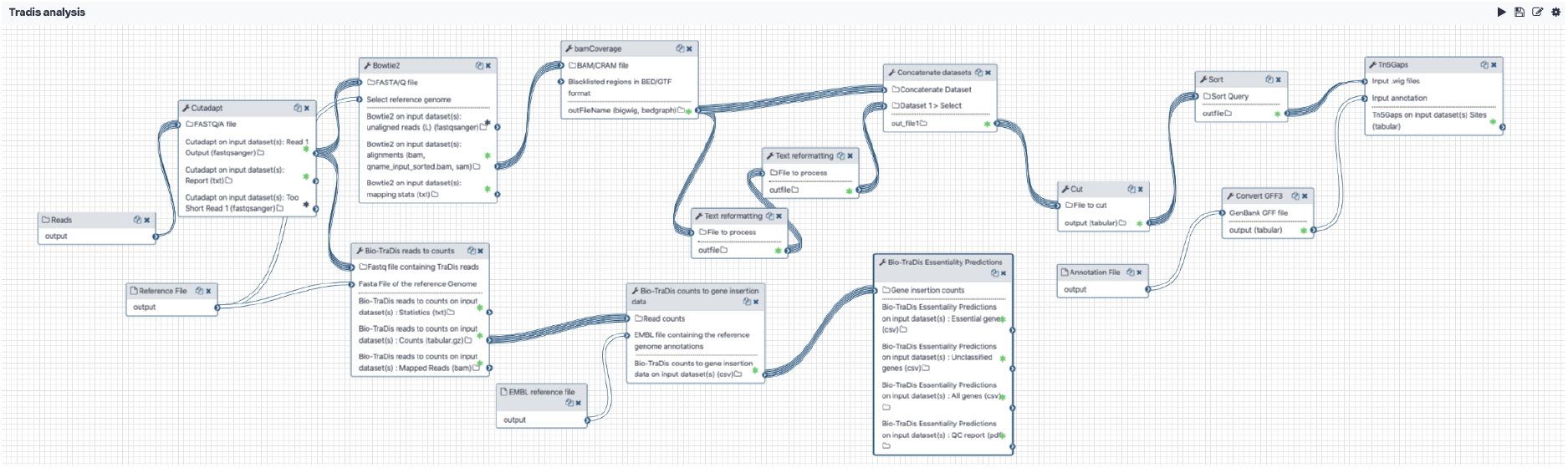
Tradis - Galaxy Workflow.

**Supplemental Figure 2.**
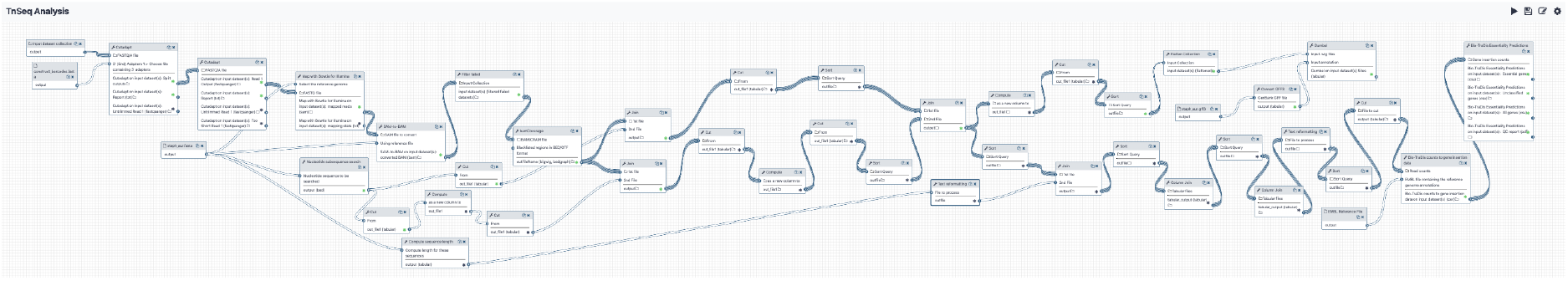
TnSeq - Galaxy Workflow.

## Informations

- Workflow for Tradis Analyses : https://usegalaxy.org/u/delphinel/w/tradis-analysis
- Workflow for TnSeq Analyses : https://usegalaxy.org/u/delphinel/w/tnseq-analysis
- History of Tradis Analyses :
- History of TnSeq Analyses :
- GitHub repository of the paper : https://github.com/galaxyproject/TIS_methods_review

